## Supplemental Figures 1-6 for "Endogenous RNAi pathway evolutionarily shapes the destiny of the antisense lncRNAs transcriptome"

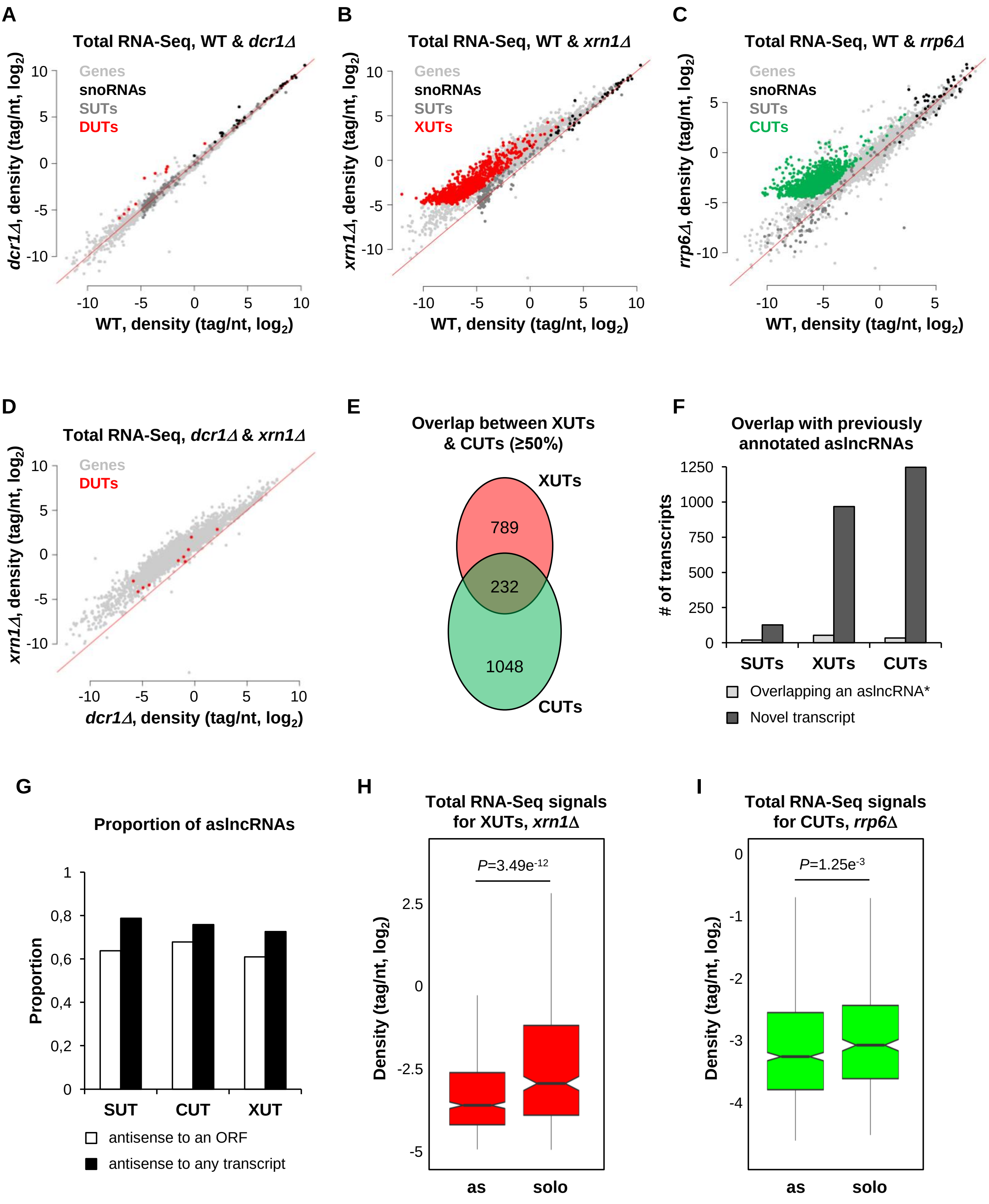
A
Total RNA-Seq, WT & dcr1Δ
Genes
snoRNAs
SUTs
DUTs
dcr1Δ, density (tag/nt, log2)
WT, density (tag/nt, log2)
B
Total RNA-Seq, WT & xrn1Δ
Genes
snoRNAs
SUTs
XUTs
xrn1Δ, density (tag/nt, log2)
WT, density (tag/nt, log2)
C
Total RNA-Seq, WT & rrp6Δ
Genes
snoRNAs
SUTs
CUTs
rrp6Δ, density (tag/nt, log2)
WT, density (tag/nt, log2)
D
Total RNA-Seq, dcr1Δ & xrn1Δ
Genes
DUTs
xrn1Δ, density (tag/nt, log2)
dcr1Δ, density (tag/nt, log2)
E
Overlap between XUTs & CUTs (≥50%)
XUTs
789
232
1048
CUTs
F
Overlap with previously annotated asIncRNAs
### of transcripts
SUTs
XUTs
CUTs
Overlapping an asIncRNA*
Novel transcript
G
Proportion of asIncRNAs
Proportion
SUT
CUT
XUT
antisense to an ORF
antisense to any transcript
H
Total RNA-Seq signals for XUTs, xrn1Δ
P=3.49e-12
Density (tag/nt, log2)
as
solo
I
Total RNA-Seq signals for CUTs, rrp6Δ
P=1.25e-3
Density (tag/nt, log2)
as
solo

**Figure S1**

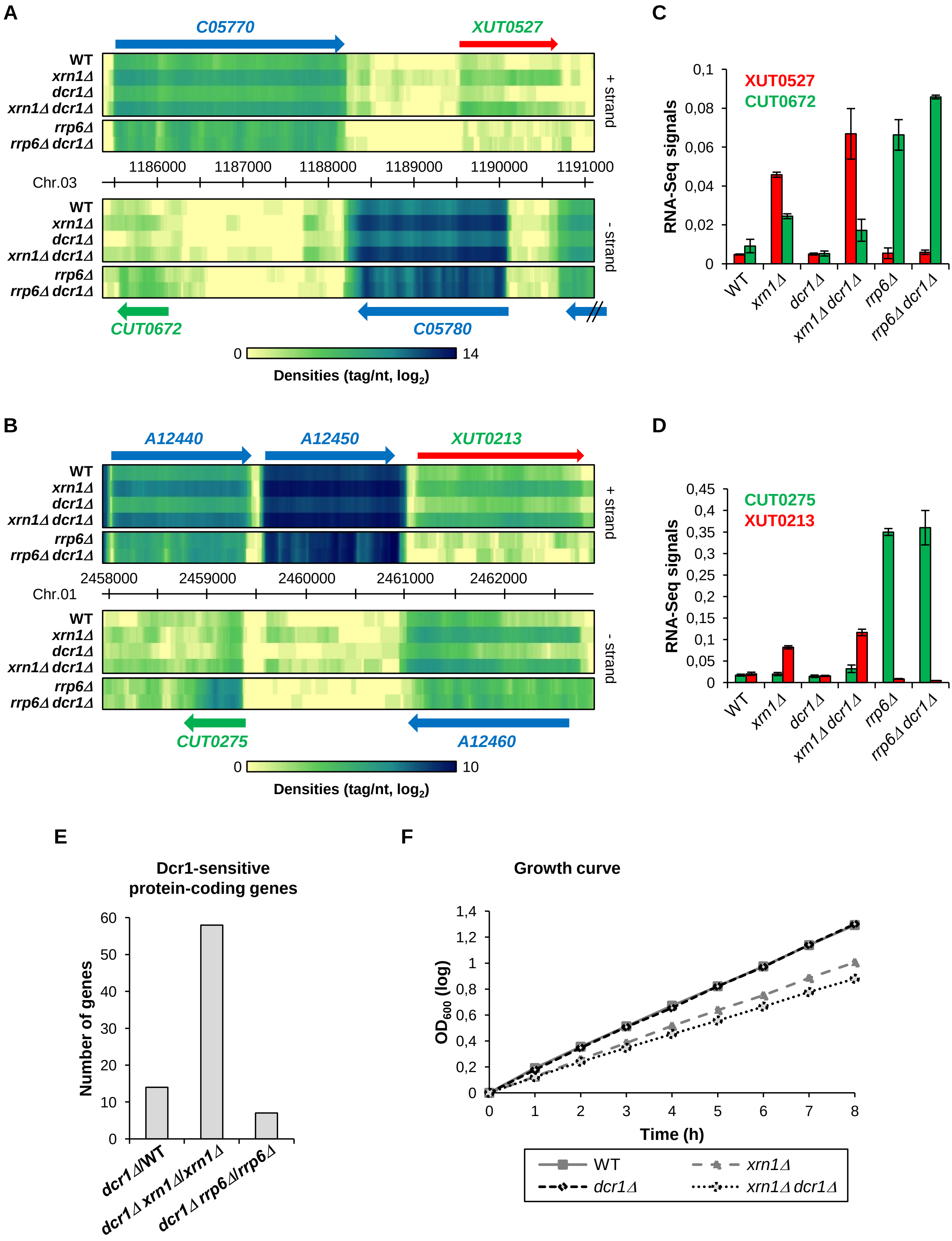
A
C05770
XUT0527
WT
xrn1Δ
dcr1Δ
xrn1Δ dcr1Δ
rrp6Δ
rrp6Δ dcr1Δ
+ strand
1186000
1187000
1188000
1189000
1190000
1191000
Chr.03
- strand
CUT0672
C05780
0
14
Densities (tag/nt, log2)
C
XUT0527
CUT0672
RNA-Seq signals
0,1
0,08
0,06
0,04
0,02
0
B
A12440
A12450
XUT0213
2458000
2459000
2460000
2461000
2462000
Chr.01
CUT0275
A12460
10
D
CUT0275
XUT0213
0,45
0,4
0,35
0,3
0,25
0,2
0,15
0,1
0,05
E
Dcr1-sensitive
protein-coding genes
Number of genes
60
50
40
30
20
10
dcr1Δ/WT
dcr1Δ xrn1Δ/xrn1Δ
dcr1Δ rrp6Δ/rrp6Δ
F
Growth curve
OD600 (log)
1,4
1,2
1
0,8
0,6
0,4
0,2
Time (h)
0
1
2
3
4
5
6
7
8
WT
xrn1Δ
dcr1Δ
xrn1Δ dcr1Δ

**Figure S2**

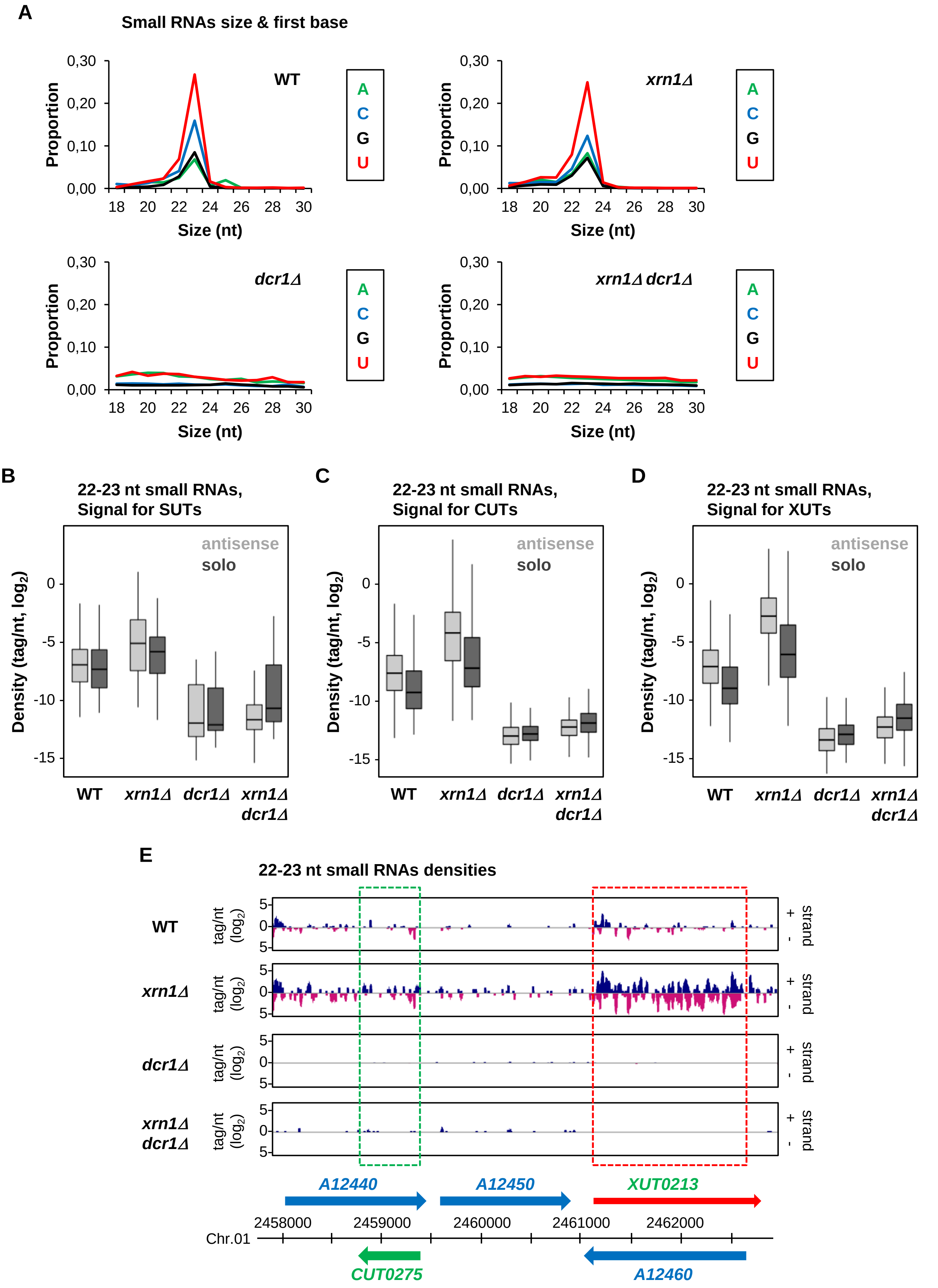
A
Small RNAs size & first base
WT
xrn1Δ
dcr1Δ
xrn1Δ dcr1Δ
A
C
G
U
Proportion
Size (nt)
B
22-23 nt small RNAs, Signal for SUTs
C
22-23 nt small RNAs, Signal for CUTs
D
22-23 nt small RNAs, Signal for XUTs
antisense
solo
Density (tag/nt, log2)
E
22-23 nt small RNAs densities
tag/nt (log2)
+ strand -
A12440
A12450
XUT0213
CUT0275
A12460
Chr.01
2458000
2459000
2460000
2461000
2462000

Figure S3

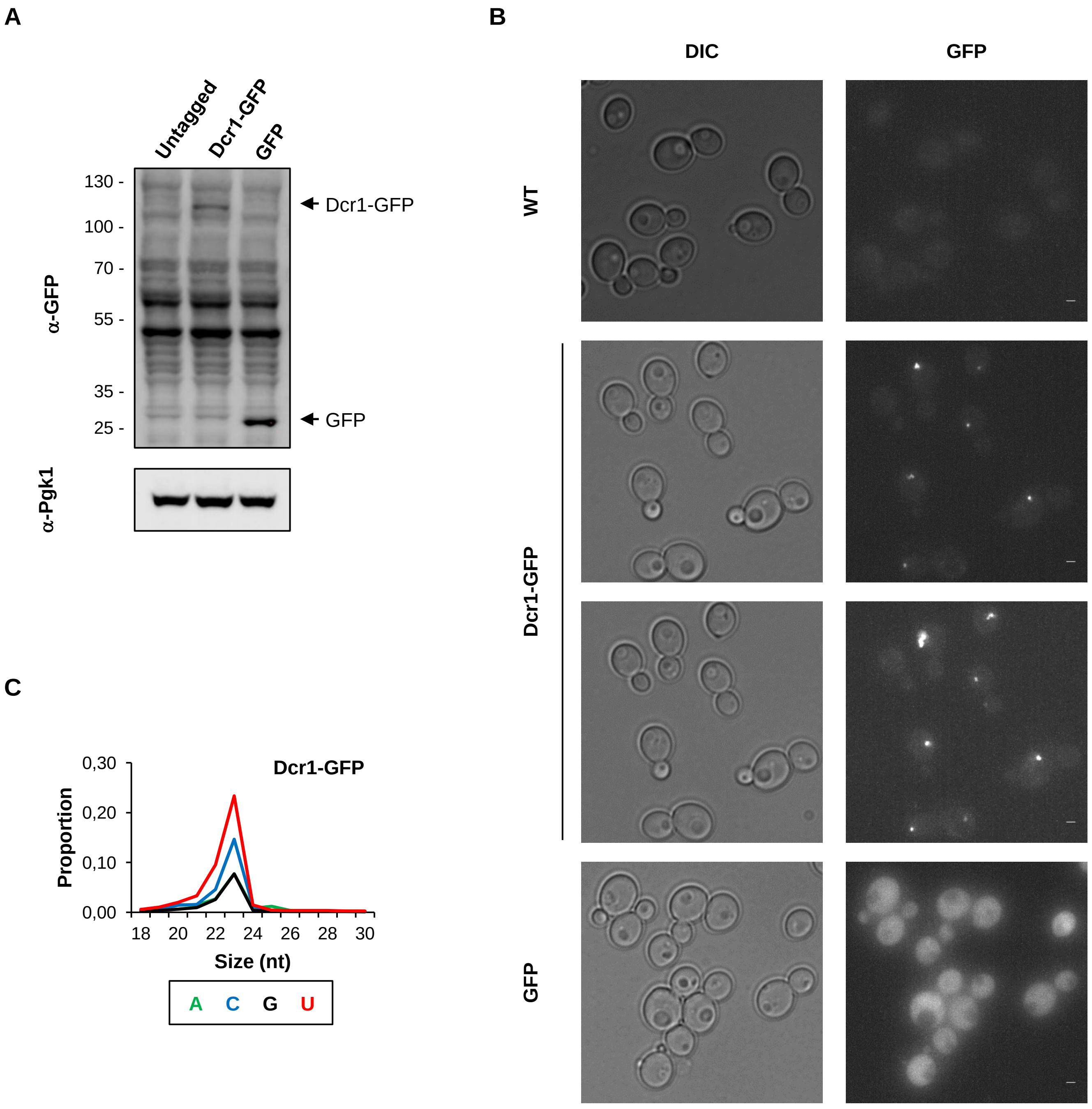
A
Untagged
Dcr1-GFP
GFP
130 -
100 -
70 -
55 -
35 -
25 -
Dcr1-GFP
GFP
α-GFP
α-Pgk1
B
DIC
GFP
WT
Dcr1-GFP
GFP
C
Dcr1-GFP
0,30
0,20
0,10
0,00
Proportion
18 20 22 24 26 28 30
Size (nt)
A C G U

**Figure S4**

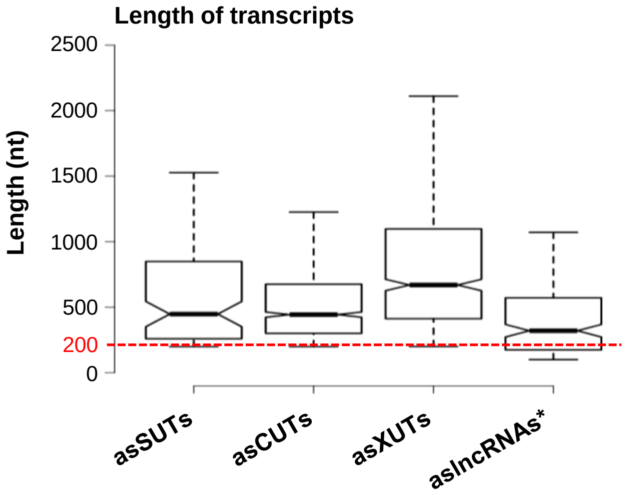
Length of transcripts
Length (nt)
2500
2000
1500
1000
500
200
0
asSUTs
asCUTs
asXUTs
aslncRNAs*

**Figure S5**

**A**

```
S.cer.   1 MGIPKFFRYISERWPMILQLIEGTQIPEFDNLYLDMNSILHNCTHGNDDDVTKRLTEEEV 60
N.cas.   1 MGIPKFFRYISERWPMILQLIEGTQIPEFDNLYLDMNSILHTCTHGNDDDVTKRMTEEEV 60
           ***************************************** ************ *****

S.cer.  61 FAKICTYIDHLFQTIKPKKIFYMAIDGVAPRAKMNQQRARRFRTAMDAEKALKKAIENGD 120
N.cas.  61 FAKIFTYIDHLFLTIKPKKTFYMAIDGVAPRAKMNQQRSRRFRTAMDAEHALQKAIDHGE 120
           **** ******* ****** ****************** ********** ** ***  *

S.cer. 121 EIPKGEPFDSNSITPGTEFMAKLTKNLQYFIHDKISNDSKWREVQIIFSGHEVPGEGEHK 180
N.cas. 121 EIPKGEPFDSNSITPGTEFMAKLTKNLKYFIHDKISNDAKWREIDIIFSGHEVPGEGEHK 180
           *************************** ********** ****  ***************

S.cer. 181 IMNFIRHLKSQKDFNQNTRHCIYGLDADLIMLGLSTHGPHFALLREEVTFGRRNSEK-KS 239
N.cas. 181 IMDFIRRITAEKDFDENTRHCIYGLDADLIILGLSTHAPHFALLREEVVFGRRNSNKVKT 240
           ** ***     ***  ************** ****** ********** ****** * *

S.cer. 240 LEHQNFYLLHLSLLREYMELEFKEIADEMQFEYNFERILDDFILVMFVIGNDFLPNLPDL 299
N.cas. 241 LENQNFYLLHLSLLREYMELEFSEIADEMQFPFDFERVLDDFILVMFVIGNDFLPNLPDL 300
           ** ******************* ********   *** **********************

S.cer. 300 HLNKGAFPVLLQTFKEALLHTDGYINEHGKINLKRLGVWLNYLSQFELLNFEKDDIDVEW 359
N.cas. 301 HLNKGAFPVILQTFKEALLHLDGYINEHGKINLERLRVWFQYLSQFELLNFEKSDIDVEW 360
           ********* ********** ************ ** **  ************ ******
```

**B**

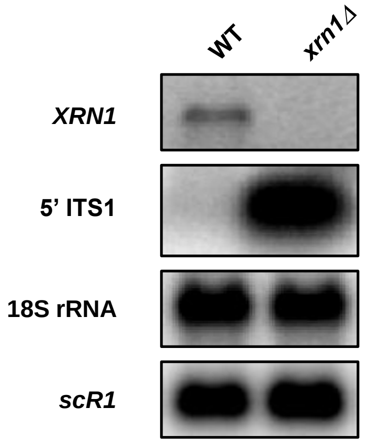


**Figure S6**
